# Tipiracil binds to uridine site and inhibits Nsp15 endoribonuclease NendoU from SARS-CoV-2

**DOI:** 10.1101/2020.06.26.173872

**Authors:** Youngchang Kim, Jacek Wower, Natalia Maltseva, Changsoo Chang, Robert Jedrzejczak, Mateusz Wilamowski, Soowon Kang, Vlad Nicolaescu, Glenn Randall, Karolina Michalska, Andrzej Joachimiak

**Affiliations:** Center for Structural Genomics of Infectious Diseases, Consortium for Advanced Science and Engineering, University of Chicago, Chicago, IL 60667, USA; Structural Biology Center, X-ray Science Division, Argonne National Laboratory, Argonne, IL 60439, USA; Auburn University, Department of Animal Sciences, Auburn, AL 36849, USA; Department of Biochemistry and Molecular Biology, University of Chicago, Chicago, IL, 60367, USA; Department of Microbiology, Ricketts Laboratory, University of Chicago, Chicago, IL, 60367, USA

**Author notes:** These authors provided equal contribution (Youngchang Kim, Jacek Wower). Correspondence should be addressed to: Andrzej Joachimiak, Structural Biology Center, X-ray Science Division, Argonne National Laboratory, Argonne, Illinois 60439, USA.

**Keywords:** Nsp15, Tipiracil, endoribonuclease, EndoU family, NendoU, SARS-CoV-2, crystal structure, COVID-19

## Abstract

SARS-CoV-2 Nsp15 is a uridylate-specific endoribonuclease with C-terminal catalytic domain belonging to the EndoU family. It degrades the polyuridine extensions in (−) sense strand of viral RNA and some non-translated RNA on (+) sense strand. This activity seems to be responsible for the interference with the innate immune response and evasion of host pattern recognition. Nsp15 is highly conserved in coronaviruses suggesting that its activity is important for virus replication. Here we report first structures with bound nucleotides and show that SARS-CoV-2 Nsp15 specifically recognizes U in a pattern previously predicted for EndoU. In the presence of manganese ions, the enzyme cleaves unpaired RNAs. Inhibitors of Nsp15 have been reported but not actively pursued into therapeutics. The current COVID-19 pandemic brought to attention the repurposing of existing drugs and the rapid identification of new antiviral compounds. Tipiracil is an FDA approved drug that is used with trifluridine in the treatment of colorectal cancer. Here, we combine crystallography, biochemical and whole cell assays, and show that this compound inhibits SARS-CoV-2 Nsp15 and interacts with the uridine binding pocket of the enzyme’s active site, providing basis for the uracil scaffold-based drug development.

## INTRODUCTION

The current pandemic of COVID-19 is caused by Severe Acute Respiratory Syndrome Coronavirus 2 (SARS-CoV-2). As a typical member of the Coronaviridae family it is spherical, enveloped, non-segmented, (+) sense RNA virus with a large ~30 kbs genome. There is no existing vaccine or proven drug against SARS-CoV-2. Since February 2020, concerted efforts have focused on characterizing its biology and developing various detection and treatment options, ranging from vaccines through antibodies to antivirals.

The coronaviruses (+) sense RNA genomes are used as mRNA for translation of a large replicase made of two large polyproteins, pp1a and pp1ab, and as a template for replication of its own copy ^1^. These polypeptides are processed by two viral proteases: papain-like protease (PLpro, encoded within Nsp3), and 3C-like protease (3CLpro or Mpro, encoded by Nsp5). For CoV-2 the cleavage yields 15 nonstructural proteins, Nsps (Nsp11 codes for just a 7-residues peptide) that assemble into a large membrane-bound replicase-transcriptase complex (RTC) and exhibits multiple enzymatic and binding activities. In addition, several sub-genomic RNAs are generated during virus proliferation from (−) sense RNA, resulting in translation of 4 structural proteins and 9-10 accessory proteins. Nonstructural proteins are potential drug targets for therapies and, because of their sequence and function conservation, developed therapeutics might in principle inhibit all human coronaviruses.

Nsp15 is a uridylate-specific endoribonuclease. Its catalytic C-terminal domain shows sequence similarity and functionality of the EndoU family enzymes, broadly distributed in viruses, archaea, bacteria, plants, humans and other animals, that are involved in RNA processing. The viral EndoU subfamily have been named NendoU. The Nsp15 enzyme is active as a 234 kDa hexamer consisting of three dimers. It was reported that NendoU cleaves both single- and double-stranded RNA at uridine sites producing 2’,3’-cyclic phosphodiester and 5’-hydroxyl termini ^2^. The 2’,3’-cyclic phosphodiester is then hydrolyzed to 3’-phosphomonoester. In coronaviruses, toroviruses and arteriviruses, NendoU is mapping to Nsp15s and Nsp11s. Nsp15s are much more sequence conserved than Nsp11s. It was proposed that in coronaviruses Nsp15 affects viral replication by interfering with the host’s innate immune response ^3,4^. To evade host pattern recognition receptor MDA5 responsible for activating the host defenses, the Nsp15 cleaves the 5′-polyuridine tracts in (−) sense viral RNAs, which are the product of polyA-templated RNA synthesis ^5^. These polyU sequences correspond to MDA5-dependent pathogen-associated molecular patterns (PAMPs) and NendoU activity limits their accumulation. For SARS-CoV it was reported that Nsp15 cleaves highly conserved non-translated RNA on (+) sense strand showing that both RNA sequence and structure are important for cleavage ^6,7^.

Recently we determined the first two crystal structures of SARS-CoV-2 Nsp15 and showed that it is similar to the SARS-CoV and MERS-CoV homologs. We have also shown that purified recombinant SARS-CoV-2 Nsp15 can efficiently cleave synthetic oligonucleotide substrate containing single rU, ^5′-6-FAM-^dArUdAdA^-6-TAMRA-3′ 8^. Studies of NendoU subfamily members, Nsp15 and Nsp11, indicate that the enzymes vary in their requirement for metal ion, with Nsp15 variants showing dependence on Mn^2+^. At the same time, the proteins are often compared to metal-independent eukaryotic RNase A, which has a completely different fold, but shares active site similarities and broad function. Specifically, it carries two catalytic histidine residues, His12 and His119, that correspond to indispensable His250 and His235 in SARS-CoV Nsp15. RNase A cleaves its substrates in a two-step reaction consisting of transphosphorylation that generates 2’,3’-cyclic phosphodiester followed by hydrolysis leading to 3’-phosphoryl terminus. The latter step has only been reported for SARS-CoV Nsp15 ^9^. Here, we have expanded Nsp15 research to explore endoribonuclease sequence specificity, metal ion dependence and catalytic mechanism. In addition to biochemical data, we report four new structures of the enzyme in complex with 5’UMP, 3’UMP, 5’GpU and Tipiracil – a uracil derivative. This compound is approved by FDA as a combination drug that is used with trifluridine in the treatment of colorectal cancer ^10^. Tipiracil inhibits the enzyme thymidine phosphorylase which metabolizes trifluridine. We discovered that Tipiracil inhibits Nsp15 endoribonuclease activity, albeit not adequately to be an effective cure, but this scaffold can serve as a template for compounds that may block virus proliferation.

## RESULTS AND DISCUSSION

### Biochemical assays

#### Nsp15 nuclease activity

We tested Nsp15 endoribonuclease activity using several assays and different substrates. We show that SARS-CoV-2 Nsp15 requires Mn^2+^ ions and retains only little activity in the presence of Mg^2+^ ions. The enzyme cleaves efficiently eicosamer 5’GAACU↓CAU↓GGACCU↓U↓GGCAG3’ at all four uridine sites (Fig. 1), as well as synthetic EndoU substrate (^5′-6-FAM-^dArU↓dAdA^-6-TAMRA-3′^) ^8^ in the presence of Mn^2+^ and the reaction rate increases with metal ion concentration. The cleavage of the eicosamer seems to show no sequence preference as 5’CU↓C, 5’AU↓G, 5’CU↓U and 5‘UU↓G are recognized and cut, especially at higher manganese concentration (Fig. 1). At 5 mM manganese, all four oligonucleotides accumulate, with the slowest cut sequence being the 5’AU↓G site. The eicosamer that does not have uridine in the sequence is not cleaved. The observed CoV-2 Nsp15 Mn^2+^ dependence is unlike its SARS-CoV homolog ^9^. Interestingly, by itself the Nsp15 can cut RNAs at any uridine site. But as a component of RTC with Nsp8 and Nsp12, Nsp15 becomes a site specific endonuclease which cuts RNA to leave short 5-10 uridine tails ^5^ and also cuts the 3’end of conserved non-coding region of RNA ^6,7^.

**Figure 1.**
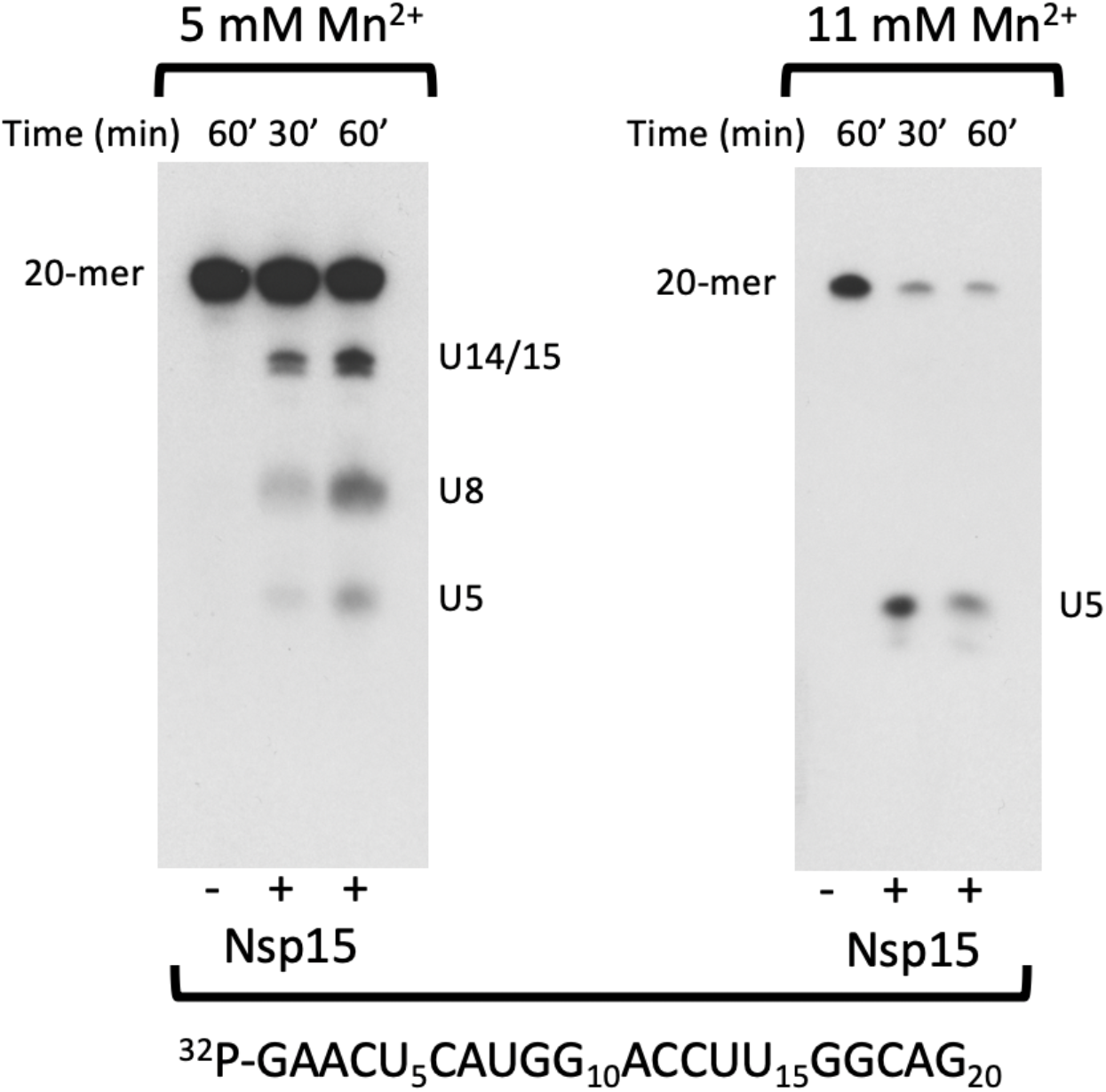
Uridine-specific endoribonuclease activity of SARS-CoV-2 Nsp15. 5’-^32^P-labeled RNA eicosamers were incubated at 37°C with Nsp15. Reaction products were separated in a 20% polyacrylamide gel containing 7 M urea.

Our observations provide new insights that might be relevant to Nsp15 roles in SARS-CoV-2 life cycle. First of all, earlier studies have demonstrated that coronavirus NendoU endonucleases that are uridine-specific display differences in their requirement for Mn^2+^ ions. On one side of the spectrum are Nsp11s of Equine Arteritis Virus (EAV), Porcine Reproductive and Respiratory Syndrome Virus (PRRSV) and Turkey Coronavirus Nsp15 that do not need Mn^2+^ for their endoribonuclease activity ^11^. SARS-Cov-2 Nsp15 might represent the other extreme end of the range. Differences in the requirement for Mn^2+^ ions may affect the outcomes of the endoribonuclease activity. For example, metal-independent Nsp11 from EAV and some Nsp15s generate oligonucleotides with 2’,3’-cyclic phosphodiesters at their 3’ ends ^9^. In contrast, metal-dependent SARS-CoV Nsp15 seems to be able to hydrolyze 2’,3’-cyclic phosphodiesters and produce oligonucleotides with 3’-phosphoryl groups on their 3’-ends ^9^. By analogy, the SARS-CoV-2 homolog should follow a two-step reaction, but this hypothesis is yet to be tested as our assays do not discriminate between the two.

The requirement for Mn^2+^ ions by SARS-CoV-2 Nsp15 may by itself enhance pathogen infectivity because Mn^2+^ is required for the host innate immune responses to viral infections. RNA binding of SARS-CoV Nsp15 was reported to be enhanced in the presence of Mn^2+^, whereas RNA binding by the cellular XendoU homolog from frog was not affected by Mn^2+^ ions ^6,7^.

The role of metal ion is puzzling as it was proposed that Nsp15 may have similar activity to eukaryotic RNase A. However, the RNase A activity is metal independent, perhaps suggesting that metal ions may help to position RNA for binding or cleavage. No metal ions or strong metal-binding sites were found in any of the models of EndoU family members available in the Protein Data Bank (PDB). Our efforts to locate Mn^2+^ ion in the Nsp15 structures also failed, arguing against a direct role of metal ions in catalysis. However, we observe the increase of Nsp15 thermal stability in the presence of such ions (Supplementary Fig. 1), though molecular basis of this behavior remains enigmatic. Previous data for SARS-CoV Nsp15, Mn^2+^ ions were found to change the intrinsic tryptophan fluorescence of SARS-CoV Nsp15, indicating conformational changes ^12,13^.

#### Inhibition by Tipiracil

Since Tipiracil is an uracil derivative and Nsp15 is specific for U, we have speculated that the compound may inhibit the enzyme. This hypothesis was tested by using heptamer RNA with single uridine site AGGAAGU, which under experimental conditions remains single-stranded and does not form duplexes. When labeled with cytidine 3’,5’-[5’-^32^P] bisphosphate (pCp), 5’-^32^P-labeled derivatives of the heptamer and unlabeled 3’CMP are produced. The transfer of the ^32^P from the pCp to the 3’ terminus of the heptamer is consistent with transphosphorylation mechanism. In the presence of 5 mM MnCl_2_, only partial cleavage is observed. This reaction is decreased by 50% in the presence of 7.5 mM Tipiracil (Fig. 2). At the higher Mn^2+^ ion concentration, the cleavage reaction proceeds to completion but no inhibition by Tipiracil can be measured (data not shown).

**Figure 2.**
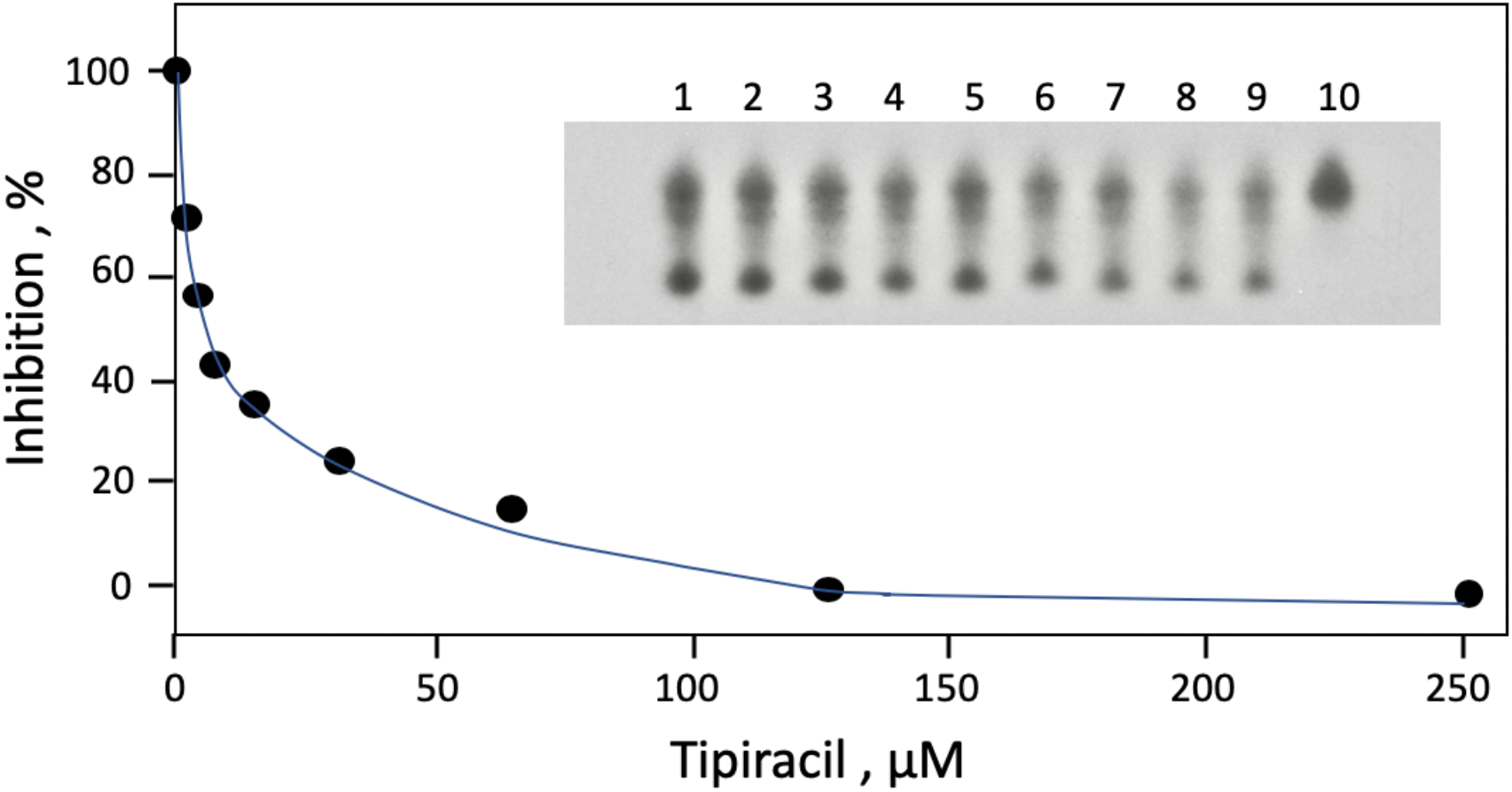
Inhibition of SARS-CoV-2 Nsp15 endoribonuclease by Tipiracil. 3’-^32^P-labeled RNA heptamers were incubated at 30°C with Nsp15 in the presence of Tipiracil at 0, 1.85, 3.9, 7.8, 15.6, 31.25, 62.5, 125.0 and 250 μM (Lanes 1-9). Lane 10: minus enzyme control. Reaction products were separated in a 20% polyacrylamide gel containing 7 M urea.

We have also performed S antigen ELISA assay in A549 cells and CoV-2 virus replication assay using qRT-PCR. Tipiracil was not affecting viability of cells but the inhibition of virus was found to be modest in the concentration range 1 – 50 mM (Fig. 3). These preliminary data suggest that the affinity of the compound may need to be improved in order to serve as an antiviral drug. Structural information is essential for this process (see below).

**Figure 3.**
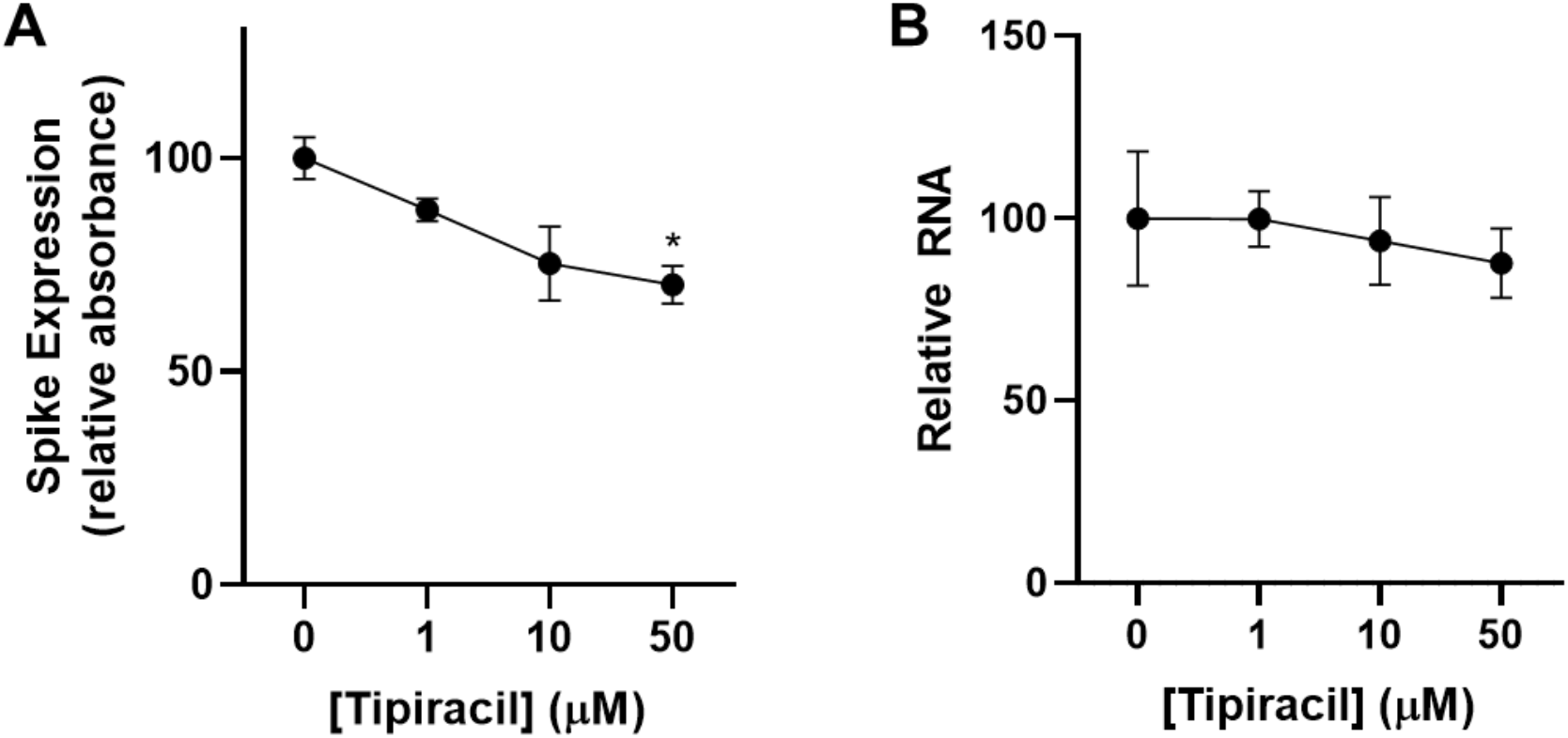
Inhibition of SARS-CoV-2 coronavirus by Tipiracil in whole cell assays. A549-hACE2 cells were pre-treated with Tipiracil or carrier (0 mM) for 2 hrs and infected with CoV-2 at MOI 1. After 48 hrs, cells were harvested to check A) spike protein and B) RNA expression. *P < 0.05.

#### Structure determination and binding of ligands

SARS-CoV-2 Nsp15 protein was crystallized with 5’UMP, 3’UMP, 5’GpU and Tipiracil using methods described previously ^8^ and the structures were determined at 1.82 Å, 1.85 Å, 1.97 Å and 1.85 Å, respectively. Crystals with ligands diffract to higher resolution than the apoprotein (2.20 Å). Of note, we were unable to obtain co-crystals of Nsp15 with 5’AMP, 3’,5’-cyclic AMP, 5’GMP, 5’TMP and 5’CMP using this same set of conditions as tested for 5’UMP. All structures were solved by molecular replacement using Nsp15 (PDB id: 6WVV) and refined as described in supplementary Materials and Methods and supplementary Table 1. The majority of the un-cleaved His-tag is not ordered and not visible but the electron density is excellent for all residues from M1 to Q347 and for all bound ligands (Fig. 4).

**Figure 4.**
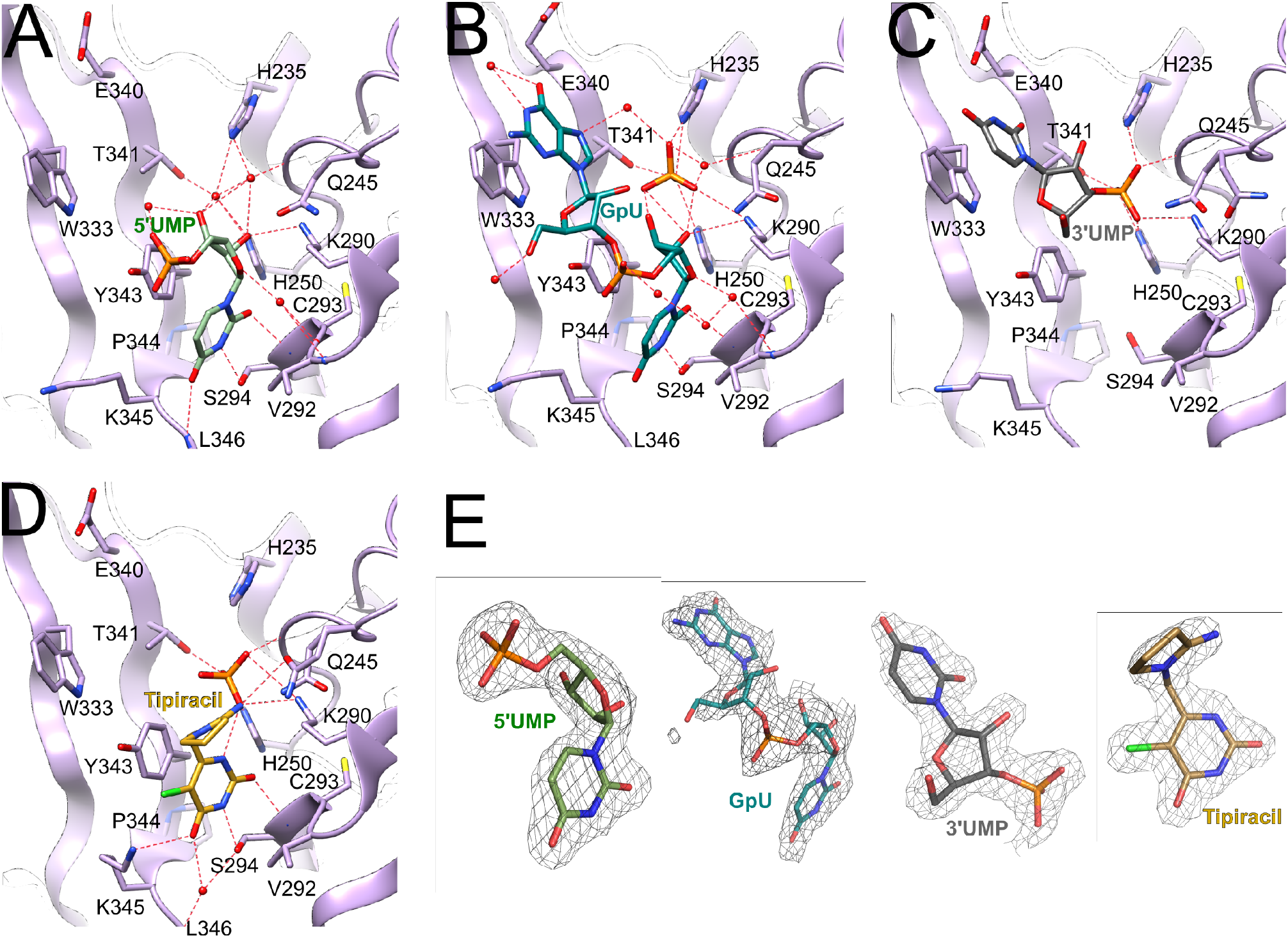
The structures of Nsp15 endoribonuclase with bound ligands. A) 5’UMP in light green, B) 5’GpU in teal, C) 3’UMP in gray and D) Tipiracil in yellow. Phosphate ions are in orange/red. E). Ligands with 2DF_o_-mF_c_ electron density maps contoured at 1.2 s.

In all four structures, ligands bind to the C-terminal catalytic domain active site (Fig. 4), though the exact positions vary, as described below. In 5’GpU and Tipiracil complexes, there is also a phosphate ion bound in the catalytic pocket. The compound binding is facilitated by side chains of seven conserved residues His235, His250, Lys290, Trp333, Thr341, Tyr343, and Ser294 and main chain of Gly248, Lys345, and Val292, as well as water molecules (Fig. 4). One non-conserved residue, Gln245, is also involved through a water-mediated interaction. The interactions do not trigger any major protein conformational changes either globally or locally. In fact, the catalytic residues (His235, His250) and other active site residues discussed below show very similar conformations in all complexes (RMSD 0. 29 Å over Ca atoms for residues His235, Gly248, His250, Lys290, Trp333, Thr341, Tyr343 and Ser294 in the pairwise superposition of complexes with the apo-structure (the highest is 0.39 Å for B chain of the Tipiracil complex and the lowest is 0.29 Å for the chain B of the 5’UMP complex)). Position of phosphate ion or 3’-phosphoryl group of 3’UMP is also preserved. Specific interactions of ligands with protein are described below.

#### 5’UMP Binding

The model of uracil binding by EndoU was proposed based on the EndoU and RNase A structures ^2^, speculating that Ser294 might be responsible for base recognition. Here we describe experimental details of such sequence discrimination. The base of 5’UMP forms van der Waals contacts with Tyr343 and several hydrogen bonds with active site residues, including side chain OG and main chain nitrogen atom of Ser294 (Fig. 4A). These interactions define O2 and N3 specificity. O4 interacts with main chain nitrogen atom of Leu346 defining the uracil specificity. Potentially, cytosine and thymine pyrimidines may also bind – an amino group in position 4 of C should be compatible with the recognition pattern and a methyl group in position 5 should not interfere with binding either as uracil position 5 is solvent accessible. In fact, recognition of C has been reported for distantly related, though with similar active site, bacterial EndoU anticodon tRNase ^14^. The ribose ring makes several hydrogen bonds with protein residues: (i) 2’OH interacts with Lys290 (NZ) and with NE2 of the catalytic His250, (ii) 3’OH makes water mediated hydrogen bonds with catalytic His235 (NE2), His250 (NE2), Thr341 (OG1) and Gly248 (main chain nitrogen atom), and (iii) O4’ is hydrogen-bonded to main chain of Val292 via a water molecule. Interestingly, the 5’-phosphoryl group projects into solvent with no interaction with protein atoms. Its only ordered interaction is with 3’OH through a water molecule. This 5’-phosphoryl group location overlaps with that of Nsp15/5’GpU complex (see below). Also worth mentioning is that the ribose in the Nsp15/5’UMP together with the phosphate ion in the Nsp15/Tipiracil mimic 2’3’-cyclic phosphodiester. The structure with 5’UMP shows how the enzyme discriminates between uracil and purine bases with Ser294 serving as the key discriminatory residue, as has been hypothesized before ^6,12,15,16^.

#### 5’GpU Binding

In RNA sequence containing uracil, such as 5’NpGpUpN3’, where N corresponds to any base, the Nsp15 cleavage would produce 5’NpGpU3’p, if transphosphorylation is followed by hydrolysis of 5’NpGpU2’3’p. In the crystal structure of Nsp15/5’GpU, the dinucleoside monophosphate binds to the active site with uracil interacting with Tyr343 and Ser294 (Fig. 4B), as seen in the Nsp15/5’UMP complex. However, the distance between O4 and main chain nitrogen of Leu346 is too long to make a good hydrogen bond. This implicates some flexibility at the protein C-terminus and suggests that the amino group in position 4 of cytosine can be accommodated, as we suggested above and reported previously ^8^. The guanine ring is stacking against Trp333 and makes two hydrogen bonds with water molecules. The absence of defined base-side chain interactions suggests lack of specificity for this site in the substrate sequence. The Nsp15/5’GpU complex binds also a phosphate ion in the active site, most likely from the crystallization buffer. The ion interacts with the protein side chains (His235, His250, Thr341, and Lys290) and uridine ribose 2’OH and 3’OH groups. It most likely mimics the binding of scissile phosphoryl group of the substrate. The backbone phosphoryl group (5’ of U) faces solvent as in the Nsp15/5’UMP complex and makes a hydrogen bond to a water molecule. This structure and the Nsp15/5’UMP complex illustrate location and specificity determinants of the uridine with a 5’-phosphoryl group. The binding of guanine in the structure identifies strong base binding site at Trp333, which, however, may not necessarily be dedicated to 5’end of the oligonucleotide (see discussion below).

#### 3’UMP Binding

We co-crystallized Nsp15 with 3’UMP, assuming that the enzyme would dock it in a manner expected for uridine monophosphate nucleotide in the contexts of larger RNA substrate, preserving the uracil specific interactions described above. Surprisingly, the uracil base is anchored by Trp333 in the guanine site observed in the Nsp15/5’GpU complex (Fig. 4C), confirming that this site can accommodate purine and pyrimidine bases. The 3’-phosphoryl group occupies the phosphate ion site created by His235, His250 and Thr341, as observed in Nsp15/5’GpU. It is surprising that uracil does not go into its dedicated site, but the result demonstrates higher affinity for the base in the Trp333 site than in uracil-recognition site, potentially governed by the strong stacking interactions with the aromatic side chain that take precedence over hydrogen bonds observed in the uracil binding of 5’UMP. The identity of the Trp333-interacting base is irrelevant, especially given that the enzyme’s substrate is most likely a larger RNA molecule.

#### Tipiracil binding – a uracil derivative and Nsp15 inhibitor

The Tipiracil molecule binds to the uracil site as observed in 5’UMP and 5’GpU (Fig. 4D and 5). The molecule makes several substrate analog-like interactions. The uracil ring stacks against Tyr341 and makes hydrogen bonds with Ser294 (interacting with O2 and N3), Lys345 (O4) and His250. The N1 atom makes a hydrogen bond with a phosphate ion and through it connects to Lys290. There are also two water mediated interactions to Ser294 and main chain carbonyl oxygen atom of Val292. Iminopyrrolidin nitrogen atom binds to Gln245, representing the only interaction unique to the ligand. Nsp15 binds Tipiracil in its active site in a manner compatible with competitive inhibition and the compound and its derivatives may serve as inhibitors of the enzyme. This structure suggests that uracil alone may have similar inhibitory properties and provides basis for the uracil scaffold-based drug development.

**Figure 5.**
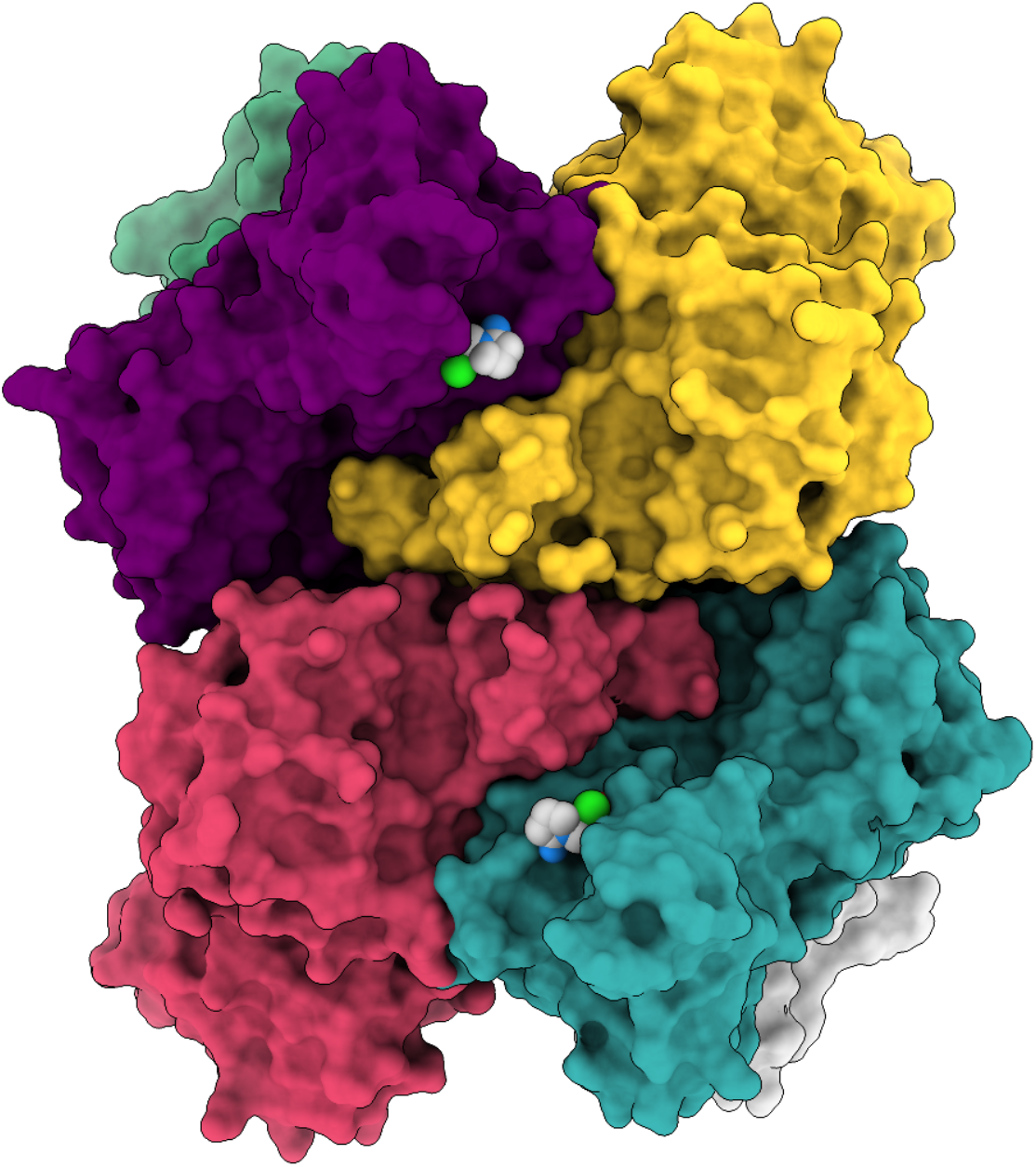
Structure of SARS-CoV-2 hexamer in surface representation showing Tipiracil bound to each subunit active site.

#### Compounds influence on Nsp15 thermal stability

The differential scanning fluorimetry experiments (DSF) showed that melting temperature (Tm) of the Nsp15 in a presence of Tipiracil, 5’GpU, 5ˈUMP, 3ˈUMP and 3’TMP is approximately 60.5°C under investigated conditions with buffer that contains 10 mM MnCl_2_ (Supplementary Fig 1. A, B). Depicted local minima of the Nsp15 first derivative of fluorescence signals have the same Tm values as a control. Although, denaturation profile of the Nsp15 in a presence of Tipiracil is broader and consistently shifted (0.5°C) to higher temperatures (Supplementary Fig 1. A). Therefore, DSF results indicates the small increase of stability of complex of the Nsp15 and Tipiracil in comparison to control sample and other tested Nsp15 complexes with 5’GpU, 5ˈUMP, 3ˈUMP and TMP. Additional change in the Nsp15 Tm is observed at 83°C (Supplementary Fig. 1B). This is caused by all ligands and may be related to increased stability of the hexamer or EndoU catalytic domain. In the presence of Mn^2+^ ions the main Tm of Nsp15 is increased from 56.5°C to 60.5°C and is Mn^2+^ concentration dependent. Interestingly, at 5 mM and 20 mM concentrations of Mn^2+^ a new local Tm minimum is observed at 83°C potentially suggesting increased stability of the EndoU domain (Supplementary Fig. 1C).

#### Comparison of CoV-2 Nsp15 active site with RNase A – mechanistic implications

Our structures of complexes with nucleotides can inform catalytic mechanism of Nsp15 endoribonuclease. We compared our structures with eukaryotic RNase A, a very well-studied model system ^17^. RNase A recognizes pyrimidine nucleotides in RNA, preferring C over U, and catalyzes a two-step reaction, the transphosphorylation of RNA to form a 2‘,3‘-cyclic CMP intermediate followed by its hydrolysis to 3‘CMP. Nsp15 also recognizes pyrimidines, preferring U over C, and was proposed to catalyze an analogous reaction ^9^. In RNase A the base selectivity is achieved by Thr45 that forms specific hydrogen bonds similar to those created by Ser294 in the Nsp15/5’UMP and Nsp15/5’GpU complexes. In RNase A the transphosphorylation reaction proceeds via an asynchronous concerted general acid/base mechanism involving His12, His119 and Lys41 ^18^. In this mechanism the 2’OH proton is transferred to the deprotonated form of His12 to activate the 2’O nucleophile. Then, the protonated His119 donates a proton to the departing 5’OH group. Lys41 function is to stabilize the negative charge that accumulates on the nonbridging phosphoryl oxygen atoms in the transition state. In the hydrolysis step the role of two histidine residues are inverted.

The RNase A active site is well organized and has several distinct pockets for binding RNA substrates (e.g. bases B1, B2, B3 and phosphoryl groups P0, P1 and P2) (for review see ^19,20^). The B1 site provides base specificity and P1 site binds the scissile phosphoester bond. B0 represent the binding site for the base upstream of the B1-P1 cleavage site. In the RNase A complex with the 5’dApdTp‾dApdA desoxyoligonucleotide (PDB id: 1RCN, ^21^), B0 interacts with adenine, B1 binds thymine, etc. The P1 site represents the above mentioned catalytic machinery consisting of His12, His119 and Lys41 and Lys41 assisted by Gln11 in phosphoryl binding.

Structural alignment of the Nsp15 and RNase A catalytic site residues and RNA ligands shows that they adopt similar positions in the two enzymes. Specifically, His250, Ser294 and Lys290 virtually overlap with the His12, Thr45 and Lys41. His250, by analogy His12 in RNase A, is in position to serve as the key residue in deprotonation of 2’OH. It is close to 2’OH in both 5’GpU and 5’UMP structures (3.2-3.8 Å respectively) and is in similar orientation in the complex of RNase A with DNA (Fig. 6). Therefore, His250 is very likely to directly activate 2’OH. Lys290 seems to play a role of Lys41 in RNase A. Main chain amide of Gly248 may provide function of Gln11 in binding to the substrate phosphoryl group. If His250 is a base in Nsp15 then the His235 must be a proton donor for the departing 5’OH group and equivalent of His119 of RNase A. However, these residues are ~8 Å apart and approach the phosphoryl group from different direction (Fig. 6A). In RNase A, His119 forms a hydrogen bond with Asp121 which may provide proton for the reaction. The structural environment for His235 is different in Nsp15. Thr341 forms a hydrogen bond with His235 and there is also Asp240 further away that makes a water mediated hydrogen bond with His235. In the hydrolysis step of converting the 2’3’-cyclic phosphate back to 2’OH and 3’-phosphoryl group the roles of histidines are reversed and now His235 must be a base deprotonating a water molecule and His250 serves as a donor of proton for the 2’OH leaving group. A different set of interactions involving Nsp15 and RNase A catalytic residues may explain why Nsp15 activity is more sensitive to low pH (data not shown) and it is expected that kinetics of the reactions will be different.

**Figure 6.**
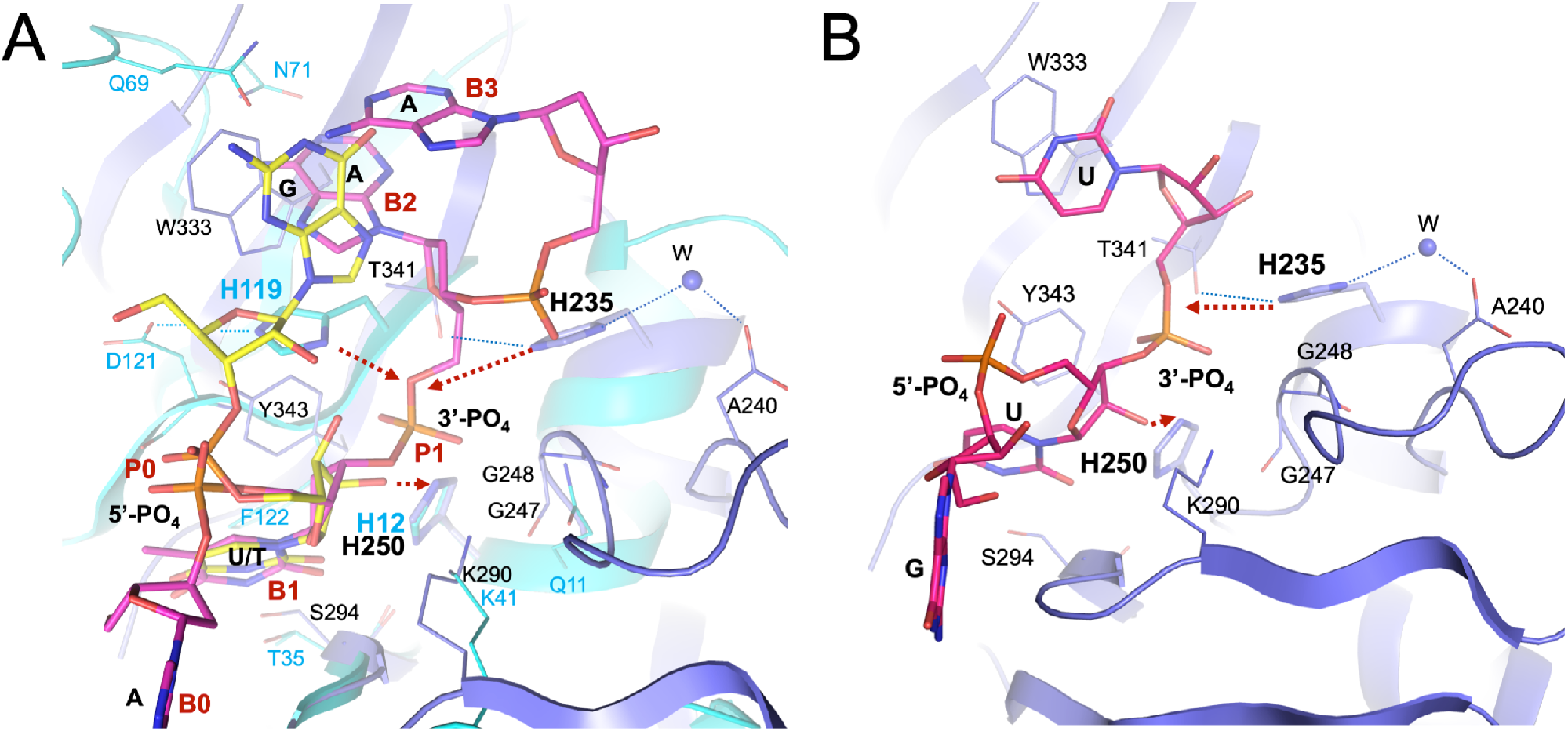
Comparison of Nsp15 and RNase A active sites. A) Nsp15/5’GpU complex compared with RNase A – 5’dApdTpdApdA complex (PDB id: 1RCN). The least square superposition is calculated with the residues H250/H12, S294/T35, K290/K41, and U/T by using coot (RMSD 0.6 Å on Ca atoms). Nsp15 is shown in dark blue, RNase A in aqua, 5’GpU in yellow and oligonucleotide in pink. Residues in Nsp15 are labeled black, RNase A residues are aqua. (B) Model of 5’GpUpU oligonucleotide binding to the Nsp15 active site. Base positions B0-B3 and phosphoryl groups positions P0 and P1 are indicated in red (bold). (B) Model of 5’GpUpU oligonucleotide binding to the Nsp15 active site. Red arrows show proton flows (extraction and donation) in A & B.

Besides similarities in P1, the two enzymes share the organization of B1 pocket. Here, Nsp15 has a very well-defined uracil recognition site made of Ser294, Tyr343 and L346 that are equivalent to Thr45, Phe120 and Ser123 in RNase A. Further extrapolation from the RNase A model of the desoxyoligonucleotide binding allows us to hypothesize that B2 site, dedicated to the base on the 3’end of scissile bond, in Nsp15 is created by Trp333. Its side chain provides stacking option for a base with no base selectivity function. Yet in our Nsp15/5’GpU structure this site is occupied by the G base located on the 5’end of U. We speculate, that for oligonucleotides that are flanked on both sides of U, mimicking the RNase A ligand, 5’ guanine (or a different base) may adopt position of 5’dA that locates to the B0 site owing to the rotation of the P-5’O bond. Then, the available Trp333 base binding site can accept base on the 3’ end of the uridine moiety, somewhat resembling our Nsp15/3’UMP structure. When we combine oligonucleotides from our structures with RNase A/DNA complex a plausible model can be built of 5’GpUp‾U nucleotide bound to the Nsp15 active site (Fig. 6B). This model underscores importance of conserved Trp333 in anchoring RNA in the active site. While Ser294 is the key residue in discriminating base, the hydrophobic interaction with Trp333 may be a significant force for ligand docking. The B3 site is not easily identifiable in the available Nsp15 structures. In addition, unlike in RNase A, where all P0-P2 subpockets contribute to the backbone binding, in Nsp15 only P1 site is currently well defined. 5’-phosphoryl groups in position P0 of the ligands do not form any direct contacts with the protein, while RNase A P0 site has Lys66 participating in RNA binding. P2 site of RNase A is created by Lys7 and it appears that His243 may fulfill such role in Nsp15.

## CONCLUSIONS

The active site of Nsp15 can accommodate all four ligands, three nucleotides (5’UMP, 3’UMP, 5’GpU) and one synthetic analog (Tipiracil) and protein interactions with these molecules seem to stabilize the active site residues and entire catalytic EndoU domain resulting in a better ordered protein. Previous mutational studies demonstrated that two histidine (His235 and His250) and one serine (Ser294) residues are essential for SARS-CoV Nsp15 activity. We now, for the first time, illustrated how Ser294 participates in uracil (or pyrimidine) recognition and we concur with previous suggestions that His250 may be key to deprotonate the 2’OH to allow nucleophilic attack on the phosphoryl group. Comparisons of Nsp15 with RNase A active site show some similarities (active site residues conservation) and indicated a common catalytic mechanism but organization of the RNA binding site is distinct, especially at sites more distant from the elements crucial for chemistry.

Our structures are consistent with the binding of single stranded nucleic acid, such as loops or bulges, as was shown for Nsp15s and other EndoUs. Both uracil and proceeding base must be in an unpaired region in order to bind to Nsp15, which is consistent with degradation of polyU tracks as reported recently ^5^. Our structures show that the proceeding base can be guanine or uracil, or other bases as well (see below). Accommodation of 5’GpU and 5’GpUpU is in agreement with previous reports of guanine preference in anterior position with respect to U and efficient degradation of polyU tracks. However, SARS-CoV-2 Nsp15 can cleave RNA, at high Mn^2+^ concentrations, at uridine sites connected to any base in anterior position; for example, the synthetic EndoU substrate used for the nuclease assay has “A”.

Nsp15 binds nucleotides to the catalytic domain of each monomer independently, therefore it is not clear why the hexamer is required for the EndoU activity of the enzyme. We show that Nsp15 can bind and hydrolyze 4, 7 and 20 nucleotide long RNA. It is possible that the hexamer is needed to bind longer RNA substrates or is involved in interactions with other proteins (for example Nsp8 and Nsp12) within RTC ^22^.

The role for metal ion requirement remains a puzzle. Although the Mn^2+^ dependence has been reported for some EndoU members and appears to be a common feature of NendoU subfamily, the metal-binding site was never located. Past studies indicated that single strand polyU RNA is relatively unstructured under most conditions. It was showed that Nsp15 cleavage is RNA sequence and structure dependent. It is possible that the metal is required for maintaining conformation of the RNA substrate during catalysis. For example, metal ions like Mg^2+^, Mn^2+^ and Zn^2+^ form complexes with purine nucleotides to affect outcome of many enzymatic reactions ^23^. Binding of Mn^2+^ ions to RNA molecules may dramatically transform their structure, as it was shown for riboswitch ^24^ and the *Bacillus subtilis* M-box aptamer that sequence contains U track (PDB id: 3PDR, ^25^). As manganese can exist in several oxidation states, it is also possible that metal redox properties can affect protein and RNA interactions and chemistry.

Tipiracil, an uracil derivative, binds to Nsp15 uracil site in a manner consistent with competitive inhibition. *In vitro* it inhibits Nsp15 RNA nuclease activity and shows modest inhibition of CoV-2 virus replication in the whole cell assay. While the compound itself is not optimal for the therapeutic applications, our work shows that uracil and its derivatives may represent a plausible starting point for nucleotide-like drug development. Moreover, interaction of Trp333 with bases may provide additional site to build much higher affinity inhibitors.

## ACKNOWLEDGEMENTS

We truthfully thank the members of the SBC at Argonne National Laboratory, especially Darren Sherrell and Alex Lavens for their help with setting beamline and data collection at beamline 19-ID. Funding for this project was provided in part by federal funds from the National Institute of Allergy and Infectious Diseases, National Institutes of Health, Department of Health and Human Services, under Contract HHSN272201700060C. The use of SBC beamlines at the Advanced Photon Source is supported by the U.S. Department of Energy (DOE) Office of Science and operated for the DOE Office of Science by Argonne National Laboratory under Contract No. DE-AC02-06CH11357. JW was supported by the Hatch program of the National Institute of Food and Agriculture, U.S. Department of Agriculture.

## SUPPLEMETAL MATERIALS

**Supplementary Table 1.**
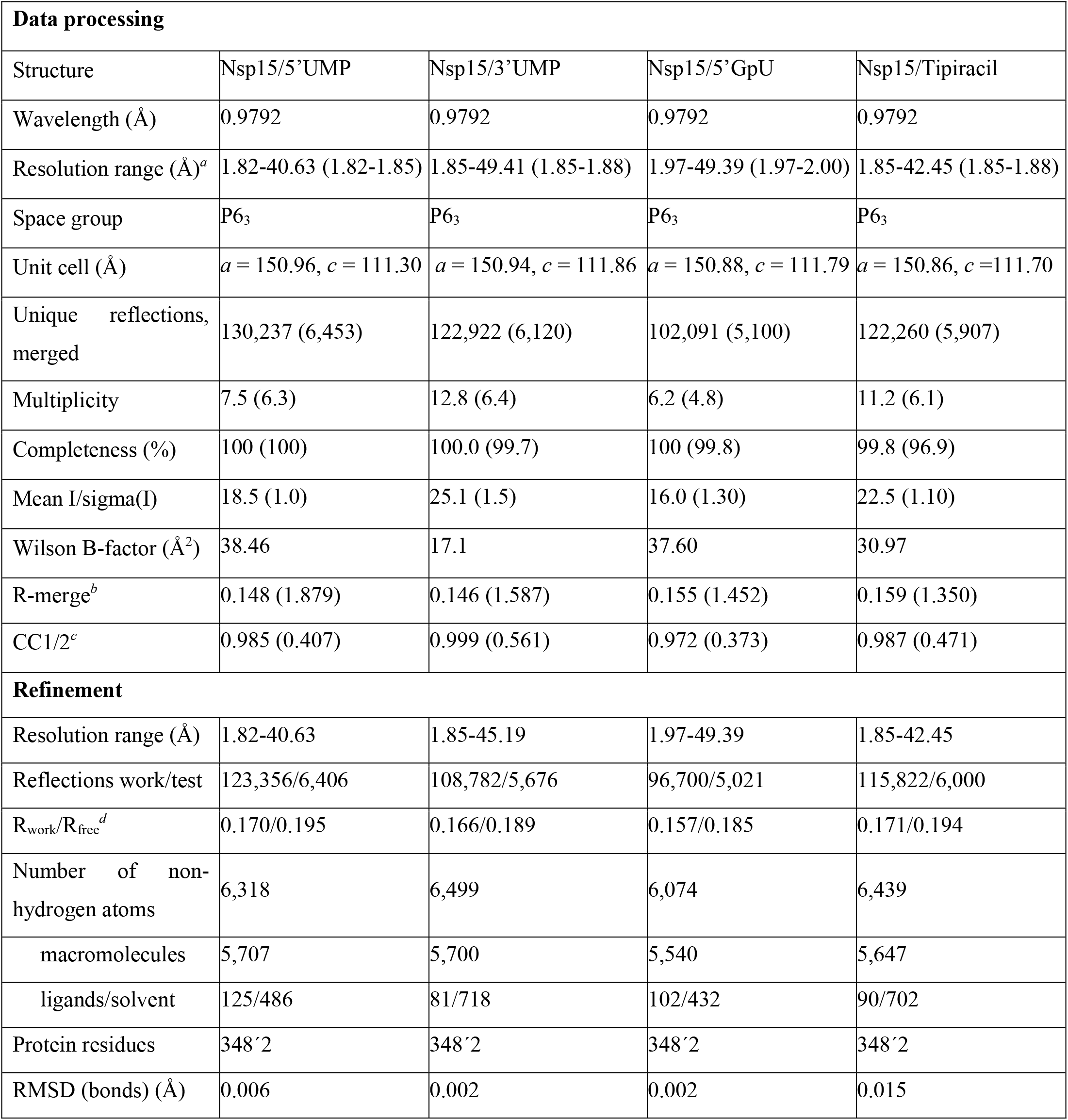
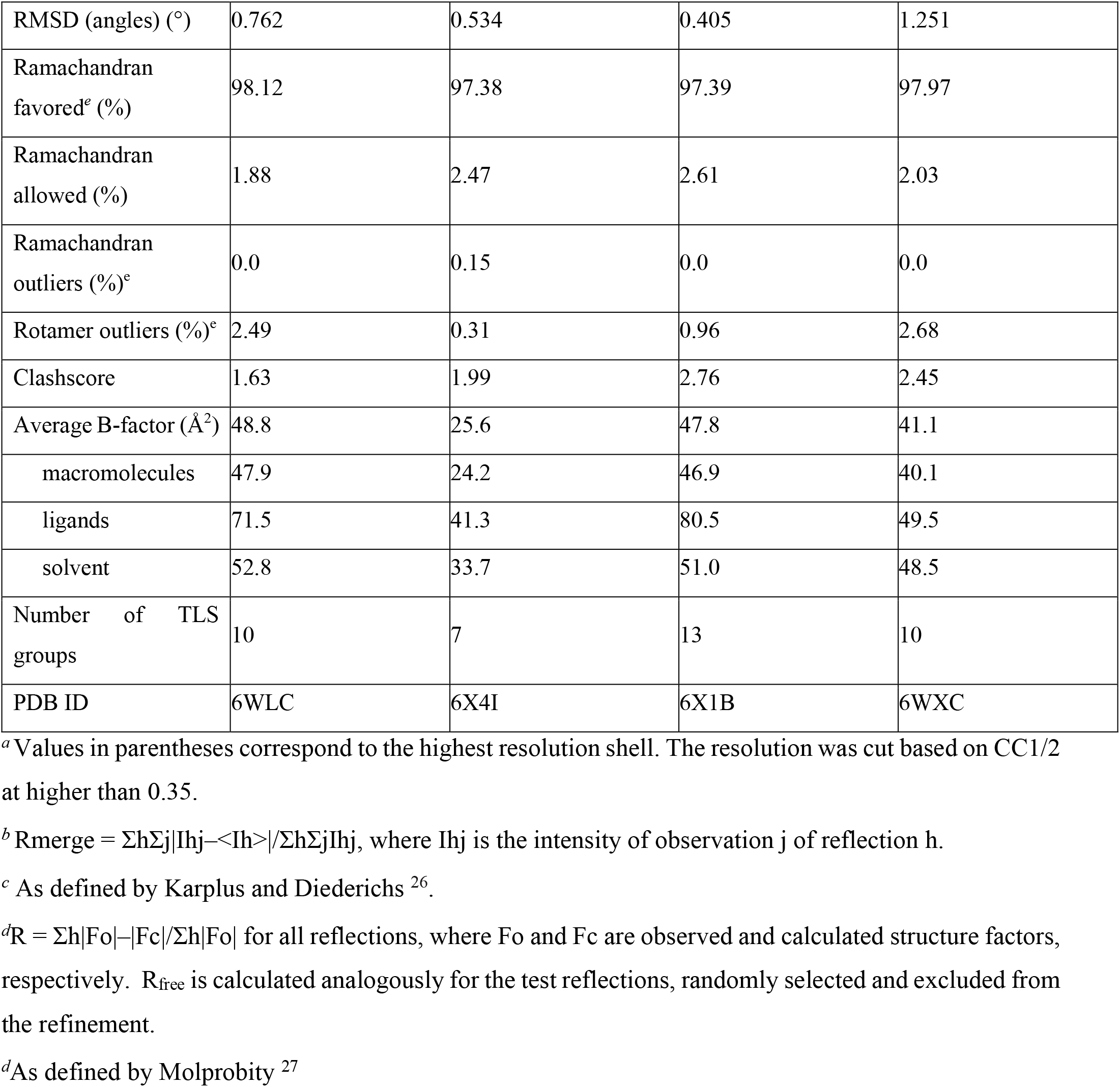
Data processing and refinement statistics.

## Supplementary Materials and Methods

3’-CMP was obtained from Sigma. Crude [gamma-^32^P]ATP was purchased from PerkinElmer. T4 Polynucleotide Kinase (3’ phosphatase minus) was from New England BioLabs. All other chemicals were reagent grade.

### Cytidine and uridine 3,5’-bisphophates synthesis

[5’-^32^P]pCp was prepared by phosphorylation of 3’-CMP as described by England et al. (1980).

### RNA synthesis

RNA eicosamers were synthetized by runoff transcription of synthetic double-stranded DNA templates as described by Sherlin et al. ^28^. 5’-^32^P-labeled RNA eicosamers and 3’-^32^P-labeled RNA heptamers were prepared according to Zwieb et al. ^29^ and England et al. ^30^, respectively.

### Protein purification and crystallization

Protein was expressed and purified as described previously. Briefly, a 4 l culture of LB Lennox medium was grown at 37°C (190 rpm) in presence of ampicillin 150 mg/ml. Once the culture reached OD_600_ ~1.0, the temperature setting was changed to 4°C. When bacterial suspension cooled down to 18°C it was supplemented with the following components to indicated concentration: 0.2 mM IPTG, 0.1% glucose, 40 mM K_2_HPO_4_. The temperature was set to 18°C for 20 h incubation. Bacterial cells were harvested by centrifugation at 7,000 *g* and cell pellets were resuspended in a 12.5 ml lysis buffer (500 mM NaCl, 5% (v/v) glycerol, 50 mM HEPES pH 8.0, 20 mM imidazole and 1 mM TCEP) per liter culture and sonicated at 120W for 5 minutes (4 sec ON, 20 sec OFF). The cellular debris was removed by centrifugation at 30,000 *g* for 1 h at 4°C. Supernatant was mixed with 4 ml of Ni^2+^ Sepharose (GE Healthcare Life Sciences) equilibrated with lysis buffer supplemented to 50 mM imidazole pH 8.0 and suspension was applied on Flex-Column (420400-2510) connected to Vac-Man vacuum manifold (Promega). Unbound proteins were washed out via controlled suction with 160 ml of lysis buffer (50 mM imidazole). Bound proteins were eluted with 20 ml of lysis buffer supplemented imidazole to 500 mM pH 8.0. Then 1 mM TCEP was added followed by Tobacco Etch Virus (TEV) protease treatment at 1:20 protease:protein ratio. The solution was left at 4°C overnight. For this particular construct, TEV protease was not able to cleave off the His tag. Nsp15 was successfully separated from TEV protease on Superdex 200 column equilibrated in lysis buffer where 10 mM β-mercaptoethanol was replaced by 1 mM TCEP. Fractions containing Nsp15 were collected. Lysis buffer was replaced on 30 kDa MWCO filter (Amicon-Millipore) via 10X concentration/dilution repeated 3 times to crystallization buffer (150 mM NaCl, 20 mM HEPES pH 7.5, 1 mM TCEP). Final concentration of Nsp15 was 19 mg/ml.

### Crystallization experiments also were conducted as described previously

All complexes, Nsp15/5’UMP, Nsp15/5’GpU, and Nsp15/Tipiracil were prepared by mixing 5-15 mM of each ligand with 0.2 mM Nsp15 and incubate for at least 30 minutes before crystallization. Crystals of Nsp15/5’GpU and Nsp15/Tipiracil, the hexagonal crystal form, in *P*63 space group, were obtained from MCSG1 screen A3 condition containing 0.2 M sodium chloride, 0.1 M sodium/potassium phosphate pH 6.2, 10 %(w/v) PEG8000. These crystals diffracted x-rays to 1.93 and 1.85 Å for Nsp15/5’GpU and Nsp15/Tipiracil, respectively. The crystals of Nsp15/5’UMP complex grew from the Crystal Screen Classic HTP (Jena bioscience) C2 condition containing 16 %(w/v) PEG4000, 0.1 M Tris pH 8.5, 0.2 M sodium acetate and diffracted to 1.82 Å. To achieve higher occupancies of the ligands in the structures, each ligand compound was added to the cryo-solution and the co-crystal was soaked for 2-3 min before frozen in the liquid nitrogen.

### Endoribonuclease Assays

Typical reaction contained 5×10^5^ CPM of 5’-^32^P-labeled RNA eicosamers or 3’-^32^P-labeled RNA heptamers (0.5 μM final RNA concentration) and 10 nM SARS-CoV-2 Nsp15 in 20 mM HEPES-KOH (pH 7.5), 50 mM KCl, 1 mM DTT and either 5 mM or 11 mM MnCl_2_. To inhibit endonuclease, Tipiciril dissolved in water was added. Following incubation at 30°C for up to 60 min, the reactions were stopped by adding an equal volume of a gel-loading buffer containing 95% formamide, 10 mM EDTA and 0.025% SDS. The products were analyzed in 20% polyacrylamide gels (acrylamide/bisacrylamide ratio, 19:1) buffered with 0.5’ Tris-borate-EDTA containing 7 M urea.

### Cells and Virus

A549 cells expressing human ACE2 (kind gift of Dr. Benjamin Tenoever, Mt. Sinai School of Medicine) were infected under biosafety level 3 conditions with SARS-CoV-2 (nCoV/Washington/1/2020, kindly provided by the National Biocontainment Laboratory, Galveston, TX).

### Cell viability

Cell viability was measured by staining with CellTracker™ Red CMTPX (ThermoFisher Scientific). CoV-2 infected cell was stained with 2 μM of CellTracker™ Red CMTPX for 30 min and then detected by Tecan Infinite m200 (Tecan) at ex577/em602 nm. After reading, cells were fixed by 10% neutral buffered formalin (NBF) for immunohistochemistry assay.

### Immunostaining against Spike protein

Immunostaining was performed on 10% NBF fixed SARS-CoV-2 infected cells in 96-well plate. After fixation, 10% NBF was removed, and cells were washed with PBS, followed by washing with PBS-T (0.1% Tween 20 in PBS), and then blocked for 30 min with PBS containing 1% BSA at room temperature. After blocking, endogenous peroxidases were quenched by 3% hydrogen peroxide for 5 min. Then, cells were washed with PBS and PBS-T and incubated with a monoclonal mouse-anti-SARS-CoV-2 spike antibody (GeneTex, 1:1000) in PBS containing 1% BSA overnight at 4°C. Primary antibody was washed with PBS and PBS-T and then cells were incubated in secondary antibody (ImmPRESS Horse Anti-Mouse IgG Polymer Reagent, Peroxidase; Vector Laboratories) for 60 min at room temperature. After washing with PBS for 10 min, color development was achieved by applying diaminobenzidine tetrahydrochloride (DAB) solution (Metal Enhanced DAB Substrate Kit; ThermoFisher Scientific) for 30 mins and detected by Tecan Infinite m200 (Tecan) at 492 nm after replacing DAB solution to PBS. Result was normalized by cell viability.

### RNA Extraction and qRT-PCR

Total RNA from SARS-CoV-2 infected cells was isolated using a NucleoSpin 96 RNA kit following the manufacturer’s instructions (Macherey-Nagel). SARS-CoV-2 RNA was quantified by qRT-PCR using SuperScript™ III Platinum™ One-Step qRT-PCR Kit w/ROX (ThermoFisher Scientific) and normalized using Eukaryotic 18S rRNA Endogenous Control (VIC™/MGB probe, Applied Biosystems) via a StepOnePlus real-time PCR system (Applied Biosystems). All reaction performed in a dual-plex qRT-PCR using the CDC recommended primers for N1. Primer and probe sequences are as follows: 2019-nCoV_N1 Forward Primer (2019-nCoV_N1-F), GACCCCAAAATCAGCGAAAT; 2019-nCoV_N1 Reverse Primer (2019-nCoV_N1-R), TCTGGTTACTGCCAGTTGAATCTG; Probe (2019-nCoV_N1-P), FAM-ACCCCGCATTACGTTTGGTGGACC-BHQ1.

### Protein differential scanning fluorimetry

The Nsp15 differential scanning fluorimetry assays were done in a buffer 20 mM Tris pH 7.5, 100 mM NaCl, 1 mM TCEP that was supplemented with SYPRO Orange (Invitrogen) dye to 5x final concentration. We used 10 μM of the Nsp15 with addition of 1 mM of Tipiracil, 5’GpU, 5ˈUMP, 3ˈUMP, TMP, respectively to get final molar ratio of protein to ligand 1:100. Additionally, the samples with ligands were supplemented with 10 mM MnCl_2_. After 30 minutes of incubation of samples at room-temperature fluorescence measurements were done using CFX Connect Real-Time System (BioRad). Fluorescent signal was detected with temperature rate 1°C per 60 sec. Graphs represents the first derivative of observed fluorescence signal. Samples were measured in triplicates and the representative standard deviation of measurements are depicted for Nsp15 control sample and complex of the Nsp15 with addition of Tipiracil.

### Data collection, structure determination and refinement

The x-ray diffraction experiments were carried out at the Structural Biology Center 19-ID beamline at the Advanced Photon Source, Argonne National Laboratory. The diffraction images were recorded at 100 K from all crystal forms on the PILATUS3 × 6M detector using 0.5° rotation and 0.5 sec exposure for 140°, 110°, and 210° for Nsp15/5’UMP, Nsp15/5’GpU and Nsp15/Tipiracil, respectively. The data were integrated and scaled with the HKL3000 suite ^31^. Intensities were converted to structure factor amplitudes in the Ctruncate program ^32,33^ from the CCP4 package ^34^ and using the apo-form SARS-CoV-2 Nsp15 structure (PDB id: 6VWW) as a search model, the structures were determined using molrep ^35^, all implemented in the HKL3000 software package. The initial solutions were refined, both rigid-body refinement and regular restrained refinement by REFMAC ^34,36^ as a part of HKL3000. The models including the ligands were manually adjusted using COOT ^37^ and then iteratively refined using COOT and PHENIX ^38^. Throughout the refinement, the same 5% of reflections were kept out throughout from the refinement (in both REFMAC and PHENIX refinement). The final structures converged to R_work_ = 0.170 and R_free_ = 0.195 for Nsp15/5’UMP, R_work_ = 0.157 and R_free_ = 0.185 for Nsp15/5’GpU and R_work_ = 0.171 and R_free_ = 0.194 for Nsp15/Tipircil with regards to each data quality. The stereochemistry of the structure was checked with PROCHECK ^39^ and the Ramachandran plot and validated with the PDB validation server. The data collection and processing statistics are given in supplemental Table 1.

## Supplementary Figures

**Supplementary Figure 1.**
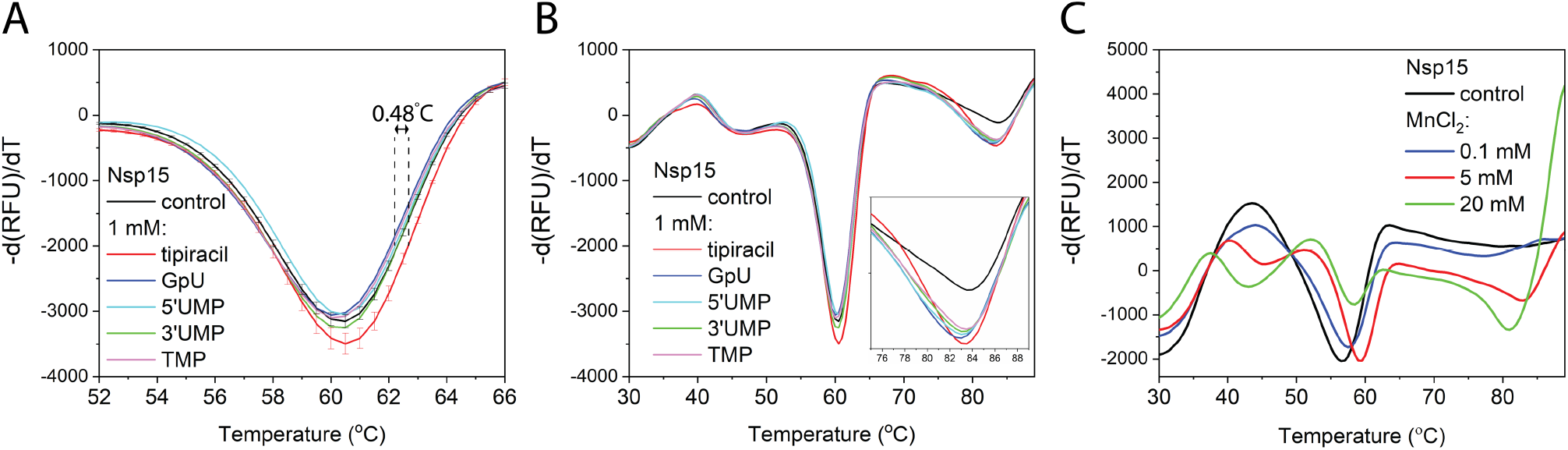
Denaturation curves of the Nsp15 in a presence of Tipiracil, 5’GpU, 5ˈUMP, 3ˈUMP, TMP (A, B). Thermal stability of Nsp15 after addition of manganese ions (C). Nsp15 samples were labeled with SYPRO orange dye ^40^.

